# β-Barrel domain swapping in α-hemolysin enables enhanced single-molecule biomolecule sensing

**DOI:** 10.64898/2026.03.12.711447

**Authors:** Chang Liu, Marco Reccia, Emilija Kavalnyte, Blasco Morozzo della Rocca, Mauro Chinappi, Jinghui Luo

## Abstract

Biological nanopores are powerful platforms for single-molecule analysis, yet rational strategies to tune their transport and sensing properties remain limited. Here we present a modular engineering approach based on β-barrel domain swapping to reprogram the function of the prototypical nanopore α-hemolysin. By replacing its native β-barrel with β-barrel domains from diverse pore-forming toxins, we generate a series of chimera nanopores that retain oligomerization capability while exhibiting reduced membrane-permeabilizing activity. Electrophysiological measurements show that selected chimera pores form stable, conductive channels with distinct ion transport properties. Notably, the αHL_NetB chimera displays stable conductance, enhanced electroosmotic flow, and improved performance in nucleic acid and protein sensing. Single-molecule experiments demonstrate that this chimera markedly slows the translocation of single-stranded DNA, enabling discrimination by length and sequence, improves the resolution of intrinsically disordered proteins such as α-synuclein, and enhances sensitivity to RNA conformational changes. Together, these results establish β-barrel domain swapping as a general and effective strategy for engineering biological nanopores with tailored single-molecule sensing capabilities.

## Introduction

Nanopore sensing is a powerful single-molecule technology that detects and characterizes biomolecules by monitoring ionic current modulations as analytes translocate through a nanoscale pore. Among available platforms, biological nanopores, typically derived from membrane proteins, offer distinct advantages over synthetic pores in terms of sensitivity, specificity and sequence-level structural definition(1). Most biological nanopores that are typically used for sensing originate from pore-forming toxins (PFTs), a diverse class of proteins produced by various pathogenic bacteria as key components of their virulence mechanisms(2). These proteins are initially synthesized as soluble monomers that, upon encountering target membranes, undergo conformational rearrangements to assemble into oligomeric complexes, which then insert into the lipid bilayer to form transmembrane pores. PFTs are broadly categorized into α-PFTs and β-PFTs, depending on whether their membrane-spanning regions consist of α-helical or β-barrel architectures(3). Based on structural and functional characteristics, PFTs can be grouped into six major families: Hemolysin, Actinoporin, ClyA, Aerolysin, Colicin, and Cholesterol-Dependent Cytolysin (CDC)(4). Among them, α-Hemolysin (αHL) from Staphylococcus aureus, has become the most widely used biological nanopore for single-molecule sensing due to its robust self-assembly, stability and well-defined structure(5,6) (Fig. 1A). Notably, αHL shares a conserved structural fold with other staphylococcal PFTs, including necrotic enteritis toxin B (NetB)(7) and *Vibrio cholerae* cytolysin (VCC)(8) (Fig. 1B, C), highlighting the modularity and evolutionary conservation of this toxin family.

**Figure 1.**
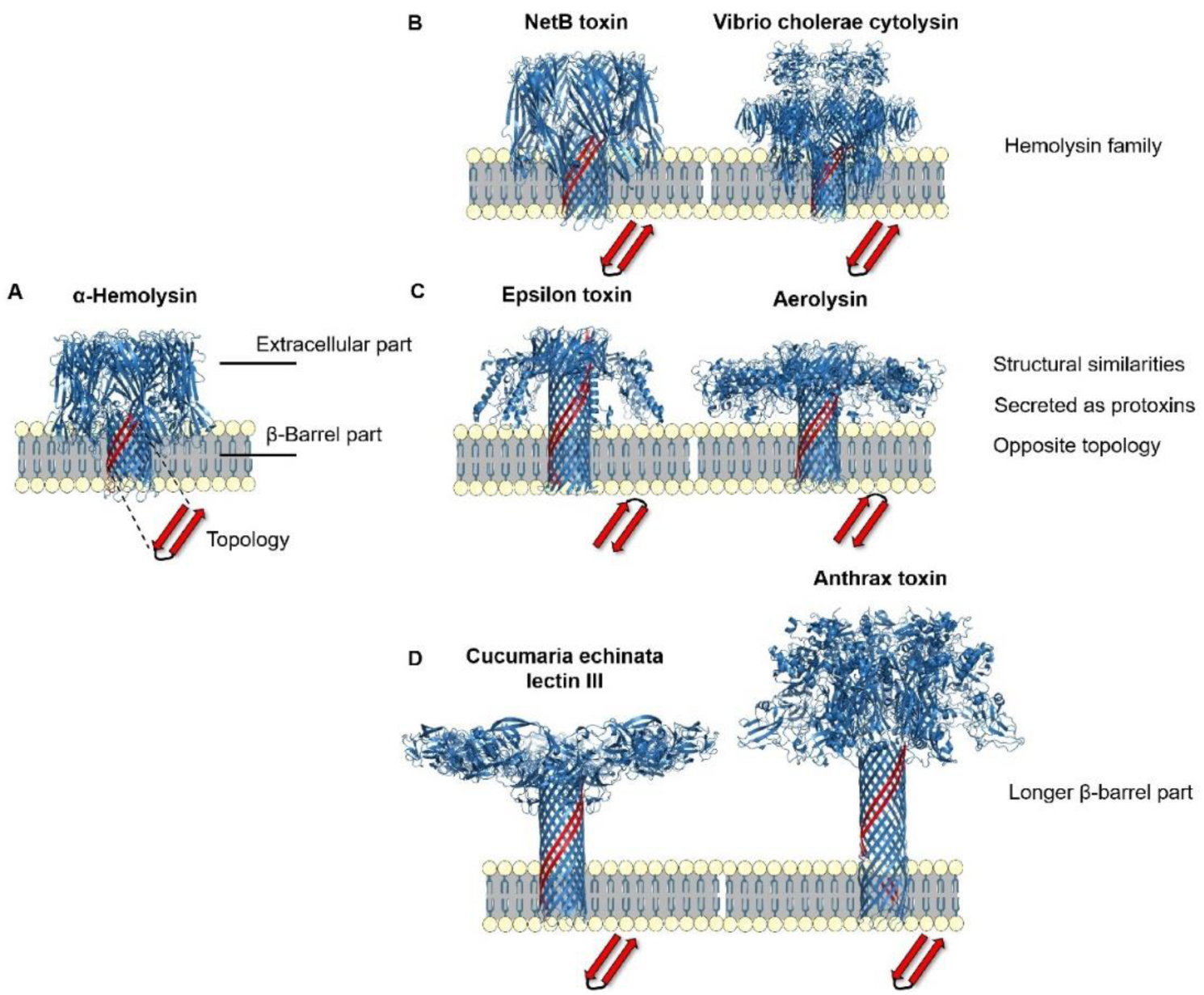
Design strategies and structural confirmation of β-PFTs. (A) α-Hemolysin (αHL) (PDB ID: 7AHL) is incorporated into a lipid bilayer (yellow and blue). The protein consists of an extracellular and a transmembrane β-barrel part. When assembled in the lipid bilayer, it shows a certain topology of β-strand, sketched as a red β-haipin. (B) Necrotic enteritis toxin B (NetB) (PDB ID: 4H56) and Vibrio cholerae cytolysin (VCC) (PDB ID: 3O44). They are from the same hemolysin family as αHL. (C) Epsilon toxin (PDB ID: 6RB9) and Aerolysin (PDB ID: 5JZT). They share structural similarities and show opposite topology compared to αHL. (D) Cucumaria echinata lectin III (CEL-III) (PDB ID: 3W9T) and Anthrax toxin (PDB ID: 6UZB).

Driven by various applications in sensing, sequencing, and molecular recognition, biological nanopore engineering has emerged as an active and rapidly evolving field(9–11). A prominent area of focus within nanopore engineering is the rational modification of existing biological nanopores to enhance their transport and sensing properties. For instance, Fragaceatoxin C (FraC) has been engineered to introduce hydrophobic gating, showcasing conductance modulation through introducing mutations in its constriction region(12). Other studies demonstrated that precise tuning of pore diameter allows discrimination of oppositely charged peptides with high mass resolution (13). In aerolysin nanopore, structure-guided point mutations guided by molecular simulations, have been used to modulate electrostatics and pore geometry, thereby enhancing their ion selectivity and biomolecular sensing capabilities (14). Further investigations include the use of protein extensions or internal adaptors to prolong analytes residence time, as demonstrated in ClyA nanopores, enabling detailed interrogation of molecular interactions (15)(16). Collectively, these advances have positioned engineered biological nanopores as promising tools for single-molecule proteomics(16), with recent studies showing that metal-ion-functionalized MspA pores can discriminate among all 20 amino acids(17). In parallel, *de novo* design of nanopores have explored the β-hairpin(18) and α-helices(19) architectures that mimic or improve upon existing natural pore functionalities. Despite these successes, point mutagenesis typically does not induce substantial structural changes in the β-barrel region of the nanopore, often preserving the overall architecture while enabling localized functional tuning. In contrast, more radical redesign approaches, like de novo pore construction, offer the potential to substantially alter pore geometry and transport behavior but often face challenges related to protein folding, membrane insertion, and structural stability(20,21). Chimera nanopores leverage known natural components, making them more reliable, stable, and easier to engineer for functional applications.

β-PFTs generally consist of two modular regions: an extracellular vestibular domain and a transmembrane β-barrel(4). These domains vary widely across toxin families in size, geometry, and charge, offering an underexplored opportunity for domain-level nanopore engineering. For example, αHL possesses a large extracellular vestibule that facilitates analyte capture and can stabilize intrinsically disordered peptides(22)(23), whereas other β-PFTs feature longer or more constricted β-barrels that can enhance sensing resolution by slowing analyte translocation (Fig. 1B and Fig. S1). We therefore hypothesized that β-barrel domain swapping could serve as a modular strategy to reprogram nanopore transport and sensing properties beyond what is achievable by point mutations.

Here, we implement this strategy by replacing the native β-barrel domain of αHL with that from other β-PFTs (Fig 1A) to generate a panel of chimera nanopores. The extracellular domain of αHL offers a stable and permissive scaffold, while the exchanged β-barrels introduce distinct transport characteristics. We designed and purified six chimera pores and systematically characterized their oligomerization, cellular membrane activity and single-channel conductance using biochemical and electrophysiological assays. Structural features were examined using AlphaFold predictions, complemented by molecular dynamics simulations of ionic current and electroosmotic flow. Finally, we demonstrate that the αHL_NetB chimera enables slowed translocation of single-stranded DNA as well as Parkinson’s α-synuclein (α-syn), and sensitive detection of RNA conformational changes using a preQ1 riboswitch aptamer, establishing β-barrel domain swapping as a general and effective strategy for engineering biological nanopores with tailored single-molecule sensing capabilities.

## Results and discussion

### Design of the chimera nanopores

Our protein engineering strategy centers on constructing chimera nanopores by swapping β-transmembrane hairpin segments from various pore-forming proteins into the scaffold of αHL, a structurally stable β-pore-forming protein (Fig. 1A). The large and accessible extracellular vestibule of αHL provides a robust scaffold that facilitates pore assembly and analyte capture, making it particularly suitable for chimera nanopore construction and single-molecule sensing.

In selecting hairpin sequences for replacement, we considered multiple biophysical parameters influencing pore performance: sequence topology, length, surface charge, and the location of restriction sites within the β-barrel region. The constriction site is typically the narrowest part of the barrel and acts as the primary sensing zone where the passage of ions and analytes is most restricted. Its geometry and electrostatic environment govern ion selectivity, analyte residence time, and signal resolution. In addition, the overall charge distribution along the pore lumen influences electroosmotic flow and electrostatic steering of analytes under an applied electric field.

We surveyed representative β-PFTs across multiple families using structural data from the Orientations of Proteins in Membranes (OPM) database. NetB and *Vibrio cholerae* cytolysin (VCC), which belong to the hemolysin family and share a membrane insertion topology similar to αHL, were selected as primary candidates for stable chimera formation (Fig. 1A, B). Both toxins exhibit negatively charged constriction regions within their β-barrels (Fig. S1A, B), suggesting favorable electrostatic environments for cation-selective transport and enhanced control of analyte translocation.

In contrast, epsilon toxin and aerolysin, although structurally related and secreted as protoxins, display an inverted transmembrane topology to αHL (Fig. 1A, C), which may complicate proper assembly when fused to the αHL extracellular domain. Nevertheless, their β-barrels exhibit relatively smooth and symmetric electrostatic profiles with extended positively charged regions (Fig. S1C, D), features that are known to promote stable analyte capture and reproducible translocation. Aerolysin, in particular, is widely used for high-resolution sensing due to its narrow β-barrel and strong electrostatic confinement(24) (14,25), which slow analyte passage and enhance signal resolution. However, its comparatively narrow extracellular domain limits analyte accessibility, motivating its combination with the more permissive αHL vestibule.

We additionally explored β-barrel domains from Cucumaria echinata lectin III (CEL-III) and anthrax toxin, both of which possess elongated transmembrane barrels (Fig. 1D). CEL-III features a centrally located constriction and a heterogeneous charge distribution along the pore wall (Fig. S1E), potentially enabling interactions with structurally diverse analytes but with reduced signal uniformity. In contrast, the anthrax toxin β-barrel contains a pronounced negatively charged constriction (Fig. S1F), which may enhance electrostatic trapping of positively charged analytes and prolong residence time within the sensing region.

Together, these considerations guided the design of a panel of chimera nanopores generated by replacing the native β-barrel of αHL with β-hairpins from structurally compatible β-PFTs. This domain-swapping strategy leverages the stability and accessibility of the αHL scaffold while introducing distinct transport and sensing properties through β-barrel exchange, providing a modular platform for tuning nanopore function and probing structure–function relationships in β-PFT assembly.

### Oligomerization and hemolysis

We assessed the assembly and pore-forming ability of six engineered chimera nanopores. SDS–PAGE analysis showed that wild-type αHL efficiently formed heptameric oligomers, whereas among the chimera constructs only αHL_VCC and αHL_NetB displayed prominent oligomeric species; the remaining designs were predominantly monomeric, indicating impaired assembly (Fig. 2A). These differences highlight the importance of structural compatibility between the αHL extracellular domain and the exchanged β-barrel.

**Figure 2.**
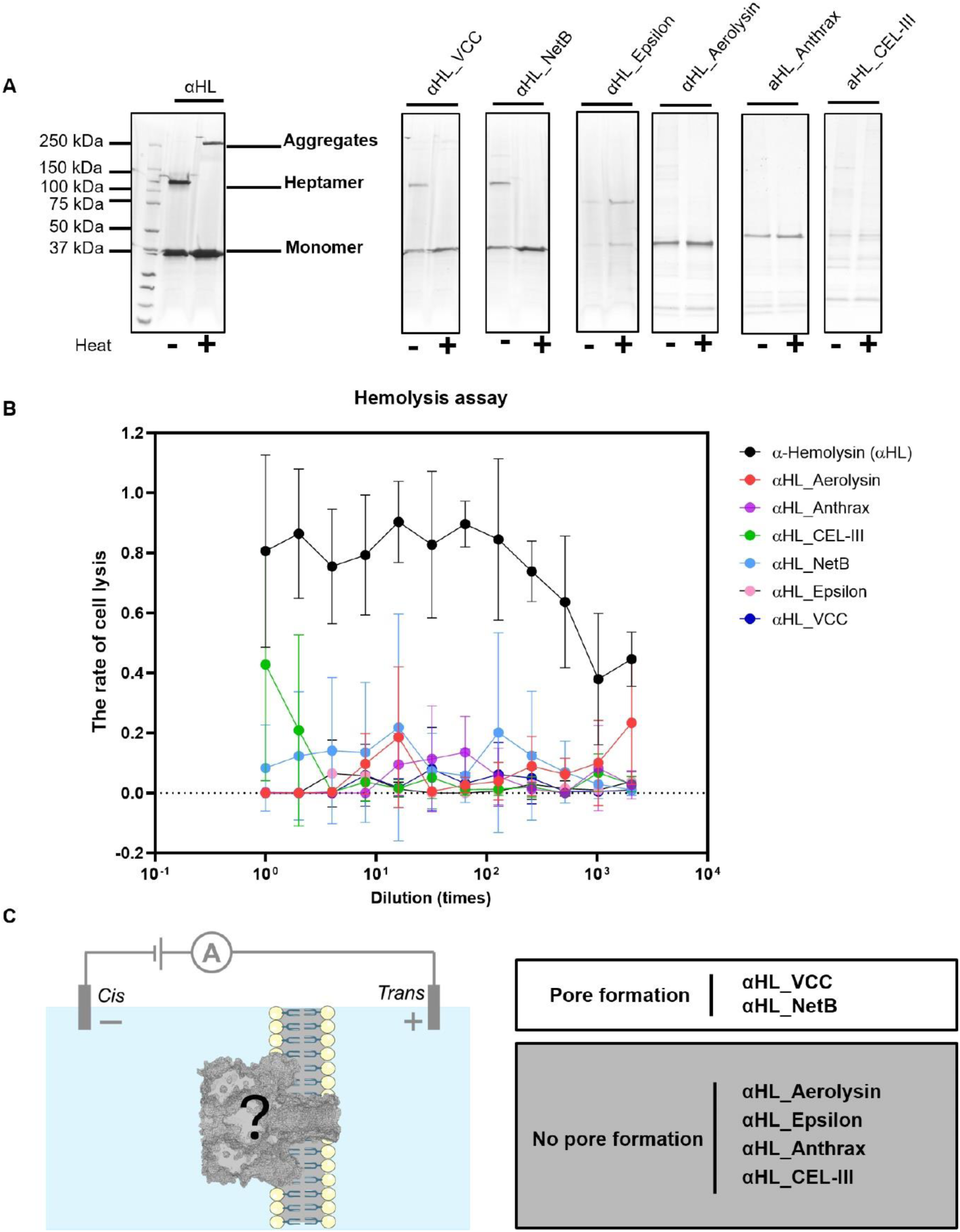
Evaluation of chimera pore assembly, hemolytic activity, and pore formation ability. (A) Oligomerization analysis of chimera pores by SDS-PAGE. Non-heated (−) and heat-denatured (+) samples were analyzed. (B) Hemolytic activity of chimera pores. The percentage of lysed red blood cells was plotted against serial dilution factors. The initial concentrations of the various toxins were as follows: αHL, 0.26 mg/mL; αHL_Aerolysin, 0.32 mg/mL; αHL_Anthrax, 0.35 mg/mL; αHL_CEL-III, 0.35 mg/mL; αHL_NetB, 0.33 mg/mL; αHL_Epsilon, 0.31 mg/mL; and αHL_VCC, 0.35 mg/mL. (C) Assessment of pore formation via single-channel recordings. Left: Schematic representation of the experimental setup for channel current measurements across a lipid bilayer. Right: Summary of pore formation outcomes. Buffer conditions: 10 mM Hepes, 1 M KCl, pH 7.4.

Consistent with these observations, wild-type αHL exhibited robust hemolytic activity, while all chimera pores showed strongly reduced or abolished hemolysis, reflecting attenuated membrane permeabilization (Fig. 2B). Single-channel electrophysiology further confirmed functional pore formation only for αHL_VCC and αHL_NetB, which produced stable, conductive ion channels across lipid bilayers; no channel activity was detected for the other chimera constructs (Fig. 2C).

Together, these results establish that efficient oligomerization is a prerequisite for functional chimera nanopore formation and identify αHL_VCC and αHL_NetB as the only designs capable of forming stable transmembrane pores, which were therefore selected for subsequent single-molecule sensing experiments.

### αHL_NetB exhibits superior stability and ion transport relative to αHL_VCC

We characterized the αHL_NetB chimera nanopore’s structure, conductance, stability, and ion transport properties. Electrostatic analysis shows that the constriction and trans entrance are negatively charged due to residues E129 and E150, indicating cation selectivity and the potential for electroosmotic flow (EOF) (Fig. 3A). This feature is critical since it suggests that the pore is cation selective and that an EOF may set in (26–29).

**Figure 3.**
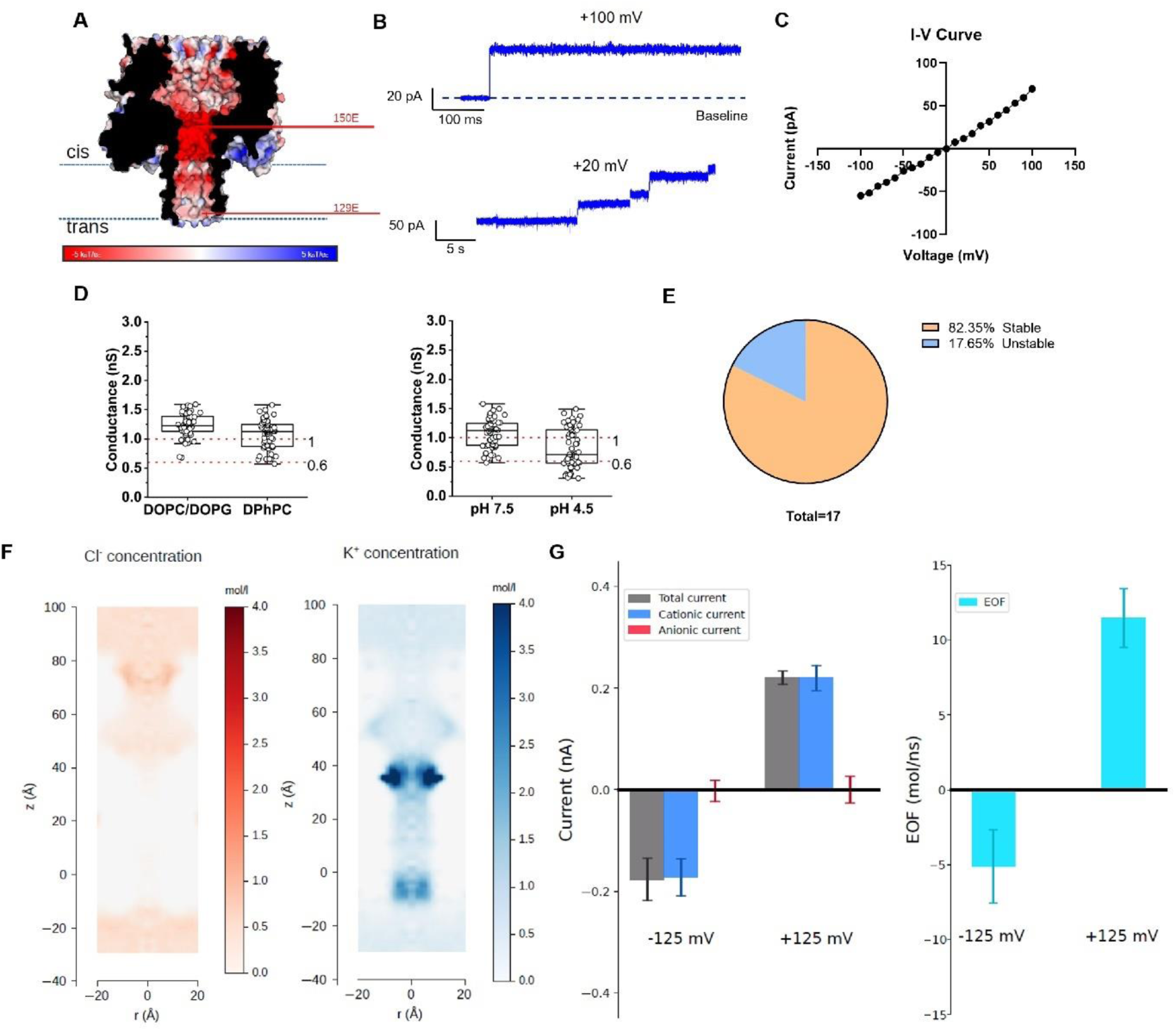
Characterization of the αHL_NetB chimera nanopore. (A) Electrostatic surface potential maps of αHL_NetB chimera pore predicted by MODELLER, computed with APBS (38), PyMOL plugin version (The PyMOL Molecular Graphics System, Version 3.1). Blue and red denote regions of positive and negative potential, respectively. (B) Representative single-channel current recordings at +100 mV and +20 mV, demonstrating stable single-pore opening and step-wise insertion. (C) I–V curve displaying linear, ohmic conductance behavior. Other I-V curves are in Figure S2. (D) Conductance measurements in different lipid compositions (DOPC: DOPG (4:1) and DPhPC) and pH conditions (pH 7.5 and pH 4.5), indicating consistent pore properties. The red dashed line indicates the conductance range observed for wild-type αHL. (E) Pore stability analysis showed that 82.35% of the pores remained stable. (F-G) MD simulations results. (F) Ion accumulation inside the pore (Cl in red, K in blue). Potassium ion accumulation and chloride ion depletion are evident, in particular in the constriction and at the trans entrance, where the residues 150E and 129E are located. (G) Cation-dominated flow and electroosmotic transport in αHL_NetB nanopore. The cationic flow (blue) is larger than anionic one (red) as expected from the negative residues exposed towards the pore. The counterions (Potassium) accumulation shown on the right panel, coupled with the external electric field, give rise to an EOF. Buffer conditions: 10 mM Hepes, 1 M KCl, pH 7.4.

Single-channel recordings at different voltages show stable and distinct open states, confirming that the αHL_NetB can properly assemble and insert into lipid bilayers to form functional pores (Fig. 3B). The nearly linear I-V curve suggests ohmic behavior without strong rectification (Fig. 3C). The conductance measurements demonstrate that the pore maintains a consistent conductance (∼1 nS) regardless of the lipid composition or pH conditions (Fig. 3D). This result suggests that the chimera pore is structurally stable and functional across different membrane environments and mildly acidic conditions, enhancing its versatility for sensing in varied experimental setups. The data also suggest that the conductance of the αHL_NetB chimera nanopore is similar to the wild-type αHL(6,27,30). Pore stability analysis shows that a significant majority (82.35%) of the pores remain stable over time, while only a small fraction exhibited instability (Fig. 3E). Fig. S3 shows the definition of stable and unstable pores. This high stability rate underlines the robustness of the αHL_NetB design, making it a promising platform for single-molecule experiments.

Ion distribution maps obtained from MD simulations reveal a notable accumulation of potassium ions near the negatively charged residues within the vestibule and pore lumen (Fig. 3F). This cation accumulation supports the hypothesis that the pore is cation selective, a hypothesis that is then validated by non-equilibrium MD simulations (Fig. 3G), where it is evident that the ionic current is dominated by cations. While the absolute conductance values derived from the MD simulations show quantitative differences when compared to the experimental data, this discrepancy is a known limitation that can be attributed to the force field parameters used in all-atom simulations (31–33). Nevertheless, non-equilibrium MD are expected to correctly reproduce the qualitative behavior of the system, providing reliable mechanistic insights on transport phenomena as selectivity and electrohydrodynamic coupling. Non-equilibrium MD simulation also shows EOF directed as the cation flow (Fig. 3G). This property is particularly beneficial for single-molecule sensing because EOF can be exploited to control the capture rate (26,34–36) and the translocation (29,37) of molecules independently on their charge. This is particularly relevant for protein sensing applications. Overall, αHL_NetB combines structural robustness, stable conductance, and EOF-driven transport, establishing it as a versatile platform for single-molecule sensing applications.

We next evaluated the αHL_VCC chimera nanopore. Electrostatic modeling revealed regions of both positive and negative potential along the β-barrel (Fig. 4A). Single-channel recordings confirmed functional pore formation, with stable open states at +50 mV and stepwise insertion at +100 mV (Fig. 4B). The I–V relationship was nearly linear, indicating ohmic behavior with approximately symmetric ion transport (Fig. 4C). Conductance measurements across different lipid compositions and pH conditions were comparable to wild-type αHL (indicated by the red dashed lines), though slight variations were observed.

**Figure 4.**
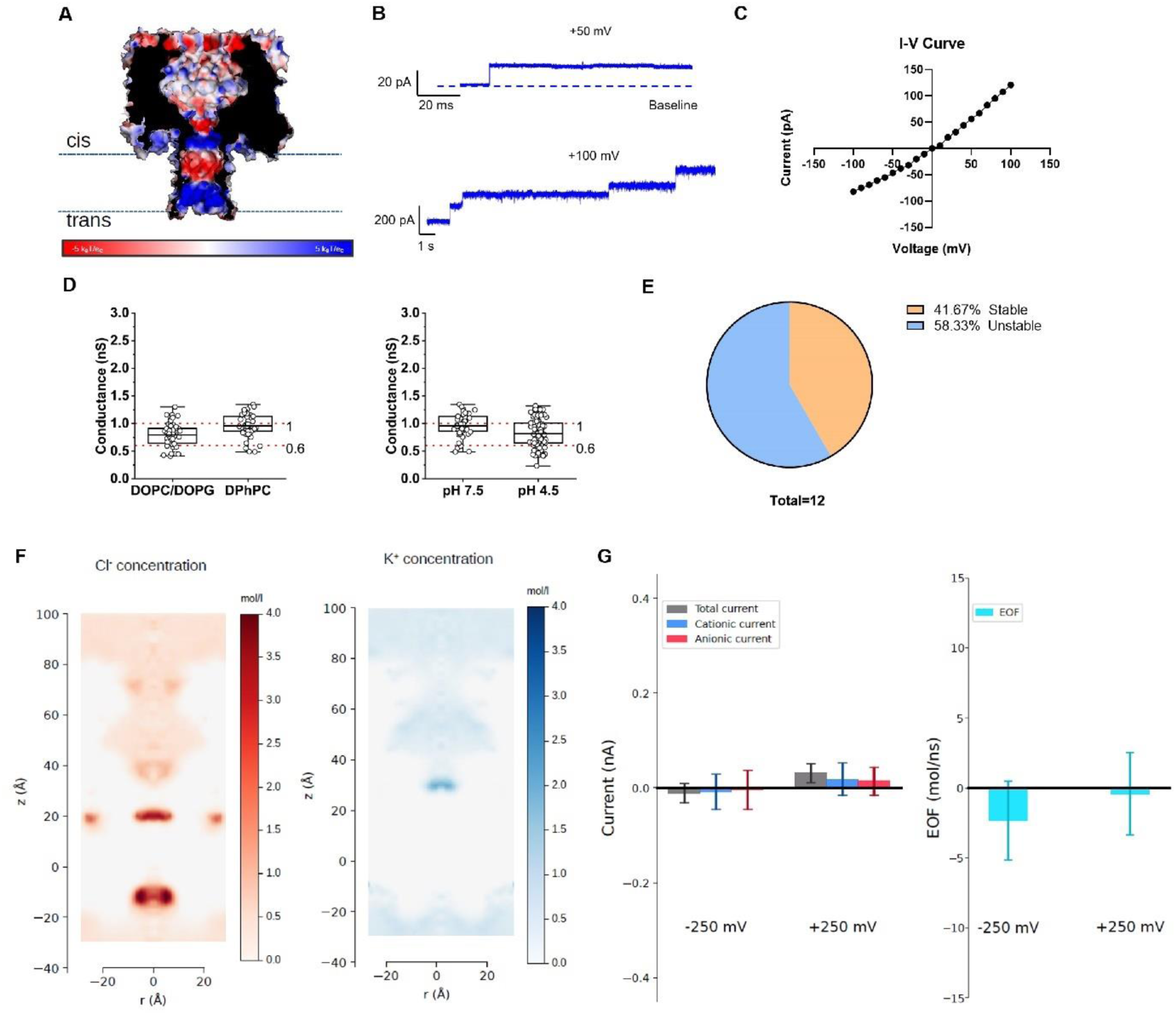
Characterization of the αHL-VCC chimera pore. (A) Electrostatic surface potential maps of αHL_VCC chimera pore predicted by MODELLER, computed with APBS (38), PyMOL plugin version (The PyMOL Molecular Graphics System, Version 3.1). Blue and red denote regions of positive and negative potential, respectively. (B) Representative single-channel recordings at +50 mV and +100 mV. (C) I–V curve showing linear conductance. (D) Single channel conductance under different lipids (DOPC: DOPG (4:1) and DPhPC) and pH conditions (pH 7.5 and pH 4.5); red dashed lines indicate wild-type αHL range. (E) Stability analysis of pores (n = 12). (F) (F-G) MD simulations results. (F) Ion accumulation inside the pore (Cl in red, K in blue). (G) The cationic flow (blue) and anionic one (red) are very low. The counterions accumulation, coupled with the geometry of the nanopore, give a very low EOF, maybe because the counterions accumulation (both positive and negative) in the constrictions creates a block for the passage of molecules (both ions than water molecules). Buffer conditions: 10 mM Hepes, 1 M KCl, pH 7.4.

Notably, the pore displays relatively high instability, with 58.33% of recordings classified as unstable (Fig. 4E), suggesting that domain swapping with VCC might compromise assembly robustness. Equilibrium MD simulation reveals a complex charge pattern in the barrel with strong accumulation of anions close to charged residues and areas with a complete depletion of cations (Fig. 4F). This is consistent with altered ion distribution caused by exposed surface charges. Finally, the non-equilibrium run indicates that no current and no EOF is observed (Fig. 4G). This is coherent with the ion distribution that, for both ions, show areas of the nanopore where there is a complete depletion of ions and that, consequently, completely block the currents. This is not coherent with experiments, where, in the stable cases, a current is observed. In our view, this is an indication that the structure predicted does not correspond to the actual experimental structure. This discrepancy is, unfortunately, not uncommon as the prediction of the structure of multimeric membrane protein is still a challenge despite the progress of AI prediction tools (39).

In contrast, αHL_NetB demonstrated higher structural stability, consistent conductance, and robust electroosmotic flow, establishing it as the preferred platform for subsequent single-molecule sensing studies.

### Detection of ssDNA using αHL_NetB chimera nanopore

Fig. 5 demonstrates the enhanced ssDNA sensing capability of the αHL_NetB chimera nanopore. The schematic fig. 5A illustrates the electrophysiological setup, where ssDNA molecules are driven through the nanopore by an applied voltage. It is observed that the αHL_NetB chimera pore significantly prolongs the duration of A40 ssDNA translocation events compared to the wild-type αHL, indicating that the chimera barrel structure effectively slows down molecular passage (Fig. 5B). Additionally, the event frequency of ssDNA translocation is reduced in the chimera pore, consistent with a more restrictive translocation process. It is shown that the αHL_NetB chimera nanopore is selective for cations (Fig. 3F). If the analyte has a different net charge than the pore-selected ion, the electrophoretic force (EP) and EOF would be antagonistic. Thus, the EOF generated by the αHL_NetB chimera nanopore can slow down the translocation speed of ssDNA.

**Figure 5.**
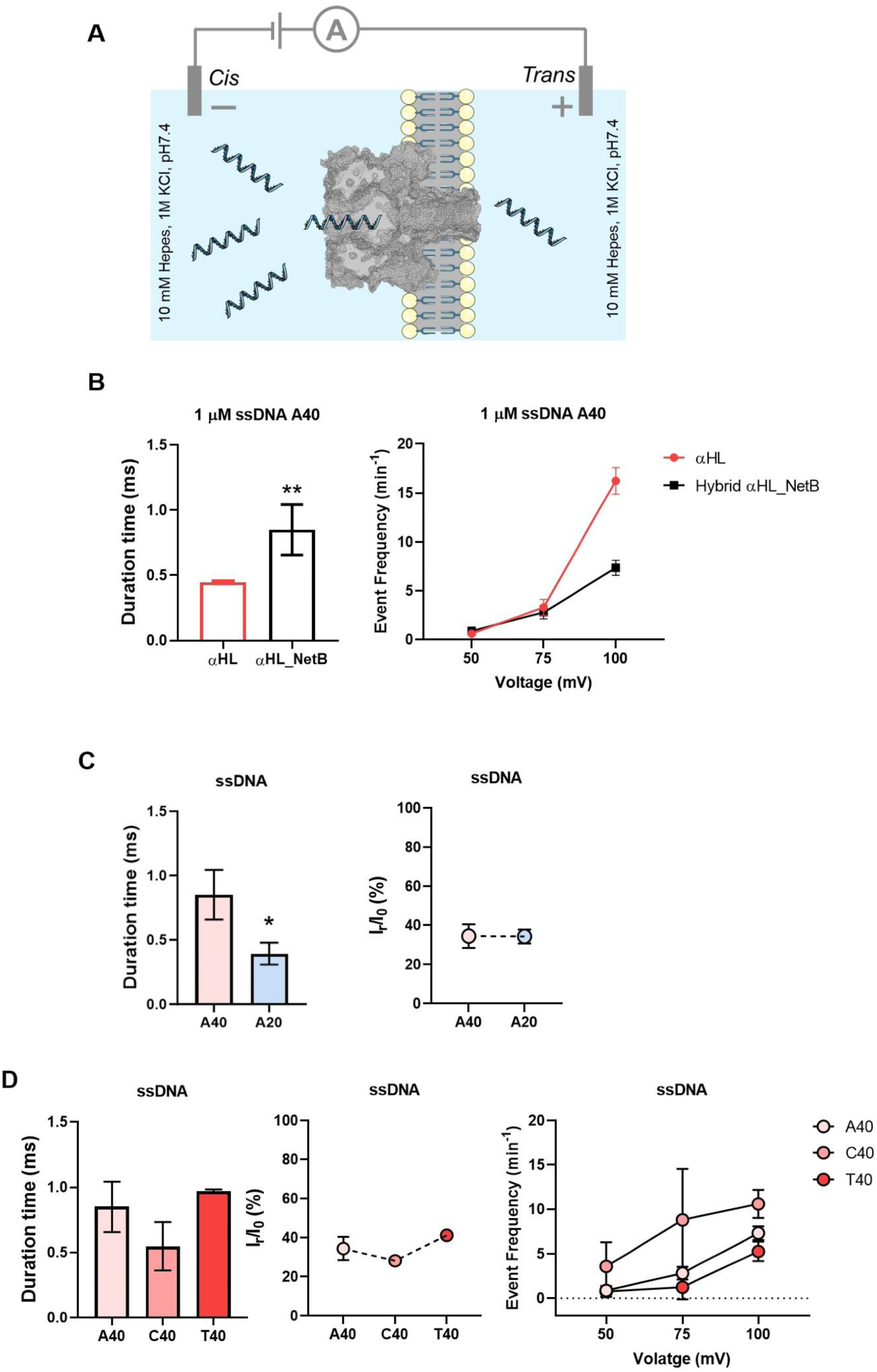
ssDNA sensing with the αHL_NetB chimera nanopore. (A) Schematic of the experimental setup for ssDNA translocation under an applied electric field across the αHL_NetB nanopore embedded in a lipid bilayer. (B) Comparison of A40 ssDNA translocation through wild-type αHL and αHL_NetB pores shows that the chimera pore significantly increases the translocation duration and decreases the event frequency. (C) Translocation measurements with A20 and A40 ssDNA indicate that longer strands exhibit prolonged dwell times, with minimal change in residual current (I_r_/I₀(%)) (*p=0.0385). (D) Discrimination of A40, C40, and T40 ssDNA strands based on translocation duration, blockade level, and event frequency, demonstrating the chimera pore’s capability for sequence- and length-dependent sensing. Error bars represent mean ± SD. Buffer conditions: 10 mM Hepes, 1 M KCl, pH 7.4.

Fig. 5C compares the translocation of A20 and A40 ssDNA strands, showing that longer DNA molecules (A40) result in increased dwell times, while the residual current (I_r_/I₀ (%)) remains similar between the two lengths, suggesting that the pore size accommodates both but senses length differences via kinetic parameters. We also examined the signals from different homopolymeric strands (A40, C40, and T40), which showed comparable patterns in dwell time, blockade amplitude, and event frequency (Fig. 5D).

Fig.S4 shows that ssDNA translocation depends strongly on voltage polarity and sequence. All homopolymers (A40, A20, C40, T40) exhibited higher event frequencies at positive voltages (50–100 mV), though the uncertainties for polyC preclude confirming a strictly monotonic increase. Event rates were negligible at negative voltages, indicating directional, voltage-driven transport. Among the sequences, polyC (C40) showed the highest translocation frequency, followed by polyA and polyT, suggesting sequence-specific differences in capture efficiency or interaction with the pore. These results highlight the importance of voltage polarity and nucleotide composition in governing ssDNA transport through the nanopore.

Overall, these results highlight the αHL_NetB chimera pore’s potential for improved nucleic acid analysis by offering enhanced control over translocation speed and enabling discrimination based on strand length and sequence composition.

### Detection of preQ1 riboswitch aptamer using αHL_NetB chimera nanopore

We next tested the αHL_NetB chimera for RNA sensing using a preQ1 riboswitch aptamer. Compared with wild-type αHL, the chimera pore substantially prolongs dwell times during aptamer translocation (Fig. 6A, B), indicating slower passage through the pore. This behavior parallels the enhanced ssDNA sensing observed in Fig. 5 and likely arises from the combined effects of the narrower effective pore and the negatively charged residues lining the β-barrel. The negative charges at the pore entrance, together with an opposing EOF, may counteract the electrophoretic force driving the negatively charged RNA, resulting in extended translocation durations. These results suggest that αHL_NetB can control RNA transport kinetics, highlighting its potential for high-resolution detection of structured nucleic acids such as riboswitches.

**Figure 6.**
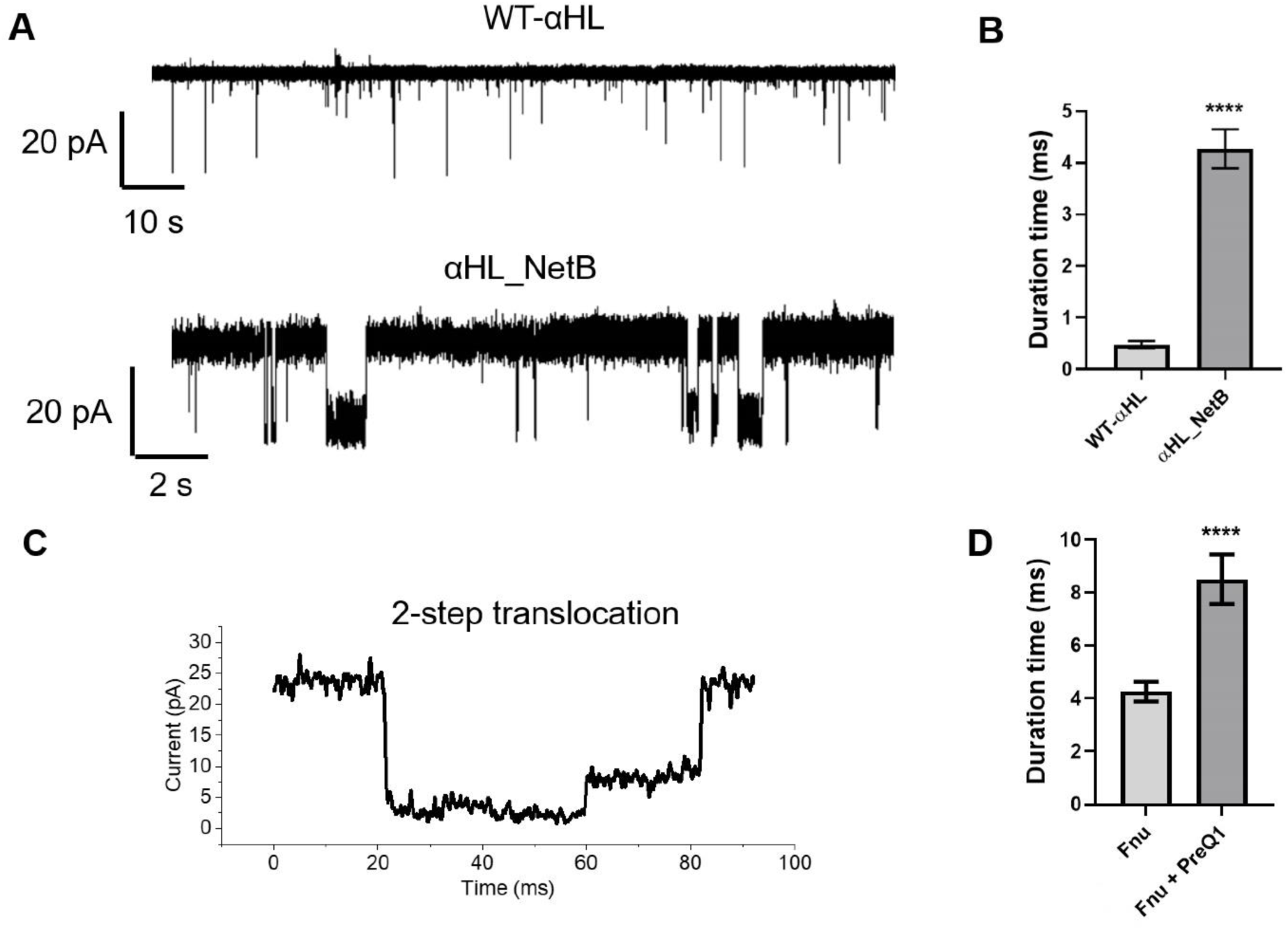
Engineered nanopores enhance preQ1 riboswitch aptamer translocation duration. (A) Representative current traces from WT-αHL and engineered αHL_NetB pores showing increased event frequency and duration with αHL_NetB. (B) Quantification of translocation durations showing significantly longer dwell times with αHL_NetB. (C) Example of a two-step preQ1 riboswitch aptamer translocation event through αHL_NetB nanopore. (D) αHL_NetB pore shows increased translocation duration of preQ1 riboswitch aptamer from *Fusobacterium nucleatum* (Fnu) in the presence of PreQ1 (****p < 0.0001). Error bars represent mean ± SEM. Buffer conditions: 10 mM Hepes, 150 mM KCl, 4 mM MgCl_2_, and pH 7.4.

This phenomenon is further supported by the results in panel D, where the addition of PreQ1(a ligand that can trigger step-wise structural change of preQ1 riboswitch aptamer from *Fusobacterium nucleatum* (Fnu), resulting in a more compact structure) also exhibits longer translocation durations through the αHL_NetB pore, suggesting that both pore structure and ligand binding can modulate translocation kinetics. Additionally, the two-step translocation event highlights the potential for these engineered systems to resolve more complex molecular features during passage (Fig. 6C). Collectively, these results demonstrate the capability of our engineered pore to enhance the resolution and sensitivity for detecting RNA conformational changes.

### Detection of α-synuclein using αHL_NetB chimera nanopore

Fig. 7 shows enhanced α-syn sensing using αHL_NetB nanopore. α-syn is a small (14 kDa) intrinsically disordered protein. Abnormal aggregates of α-syn have been implicated in various neurodegenerative diseases known as α-synucleinopathies, including Parkinson’s disease (PD), Lewy body dementia (LBD), Multiple System Atrophy (MSA), Pure autonomic failure (PAF) and REM sleep behavior disorder (RBD) (40). Consistent with earlier findings (41), adding α-syn to the cis-side of the WT-αHL pore results in almost no capture events at nanomolar concentrations. However, the αHL_NetB pore, having increased opposing EOF, enhances α-syn capture (Fig. 7B), likely due to a slowed-down drift velocity and increased hovering of α-syn close to the entrance of the pore. Such trapping of α-syn might assist in aligning the intrinsically disordered α-syn chain for threading, thereby increasing the probability of the protein to translocate rather than diffuse away from the pore. Moreover, similarly to ssDNA and preQ1 analytes, generated EOF in αHL_NetB reduces the passage duration of α-syn through the pore, with no effect on residual current (I_r_/I₀ (%)) (Fig. 7A, B). Overall, these findings suggest that engineered αHL_NetB pore improves sensing of intrinsically disordered α-syn monomers by facilitating their capture and slowing down translocation through the pore.

**Figure 7.**
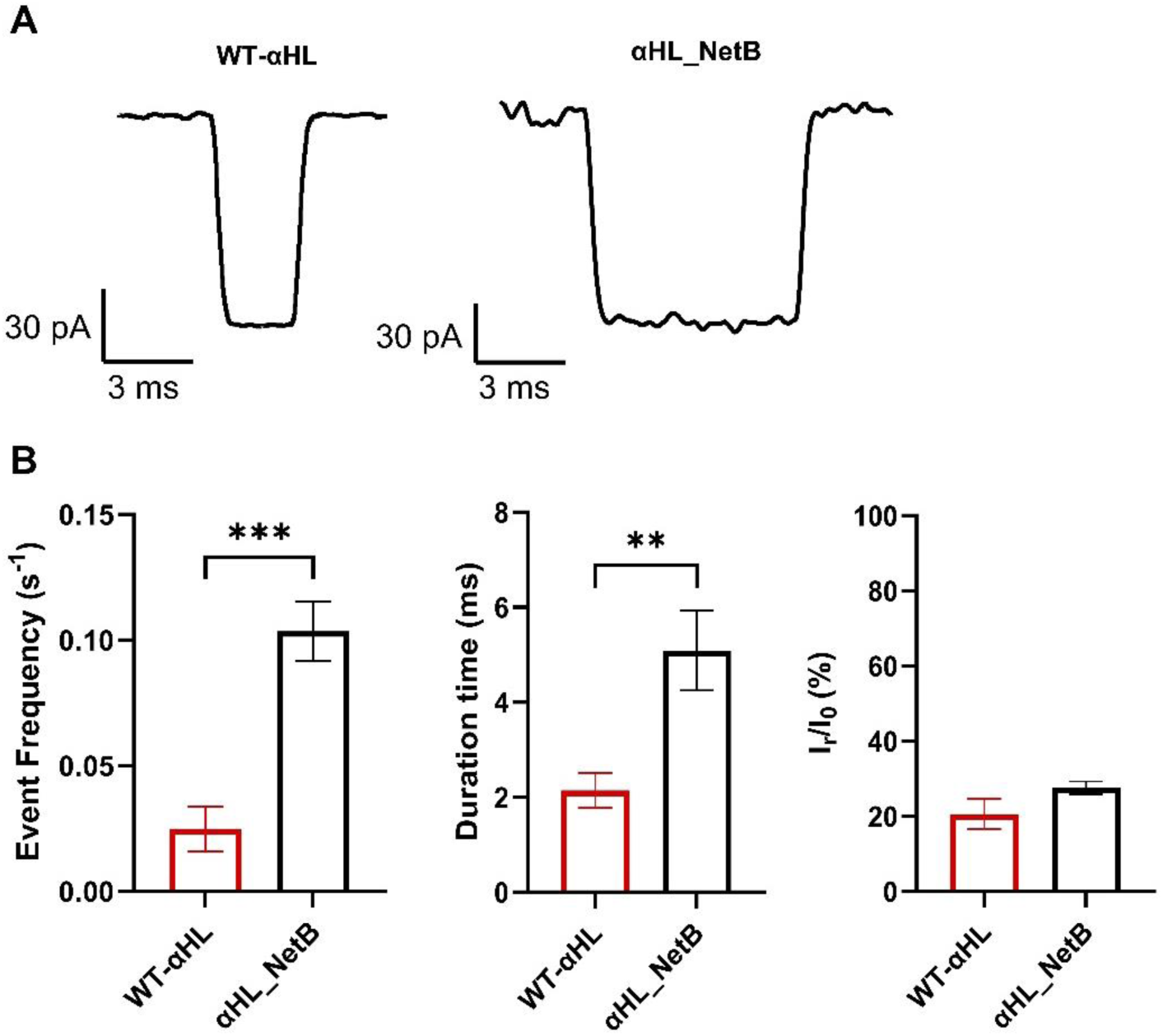
Sensing α-syn using the αHL_NetB nanopore. (A) Examples of single-molecule α-syn translocation events through WT-αHL and αHL_NetB pores. (B) Comparison of α-syn translocation between the WT-αHL and αHL_NetB pores indicating an increased event frequency and duration time for αHL_NetB, with no change in residual current (Ir/I₀ (%)) (**p = 0.002, ***p < 0.001). Error bars represent mean ± SEM. Buffer conditions: 10 mM HEPES, 1 M KCl, pH 7.4.

## Conclusions

The successful design and reconstitution of six αHL-based chimera nanopores underscore the versatility of β-barrel domain swapping as a powerful strategy for nanopore engineering. Biochemical analyses confirmed that most of the chimera pores could efficiently oligomerize but retain reduced membrane-permeabilizing activity, while electrophysiological measurements demonstrated stable channel formation with distinct conductance profiles of αHL_NetB and αHL_VCC chimera nanopores. These results not only validate the structural compatibility between αHL’s extracellular domain and diverse β-barrel domains but also establish a modular framework for creating customized nanopores with enhanced or novel sensing capabilities, advancing the development of next-generation single-molecule detection technologies. Comparative analysis highlights the critical role of β-barrel selection in defining nanopore performance. αHL_NetB exhibits high structural stability, consistent conductance, and strong electroosmotic flow, facilitating control over analyte translocation speed and enhancing single-molecule resolution for DNA, RNA, and peptides. By contrast, αHL_VCC shows reduced stability, disrupted ion transport, and minimal EOF, underscoring the importance of domain compatibility for reliable pore function.

Single-molecule experiments further establish that αHL_NetB extends ssDNA translocation durations, enabling discrimination by strand length and sequence. For neutral or weakly charged analytes, EOF dominates analyte-pore interactions, providing a tunable mechanism to enhance detection sensitivity. Similarly, the chimera pore slows preQ1 riboswitch aptamer translocation, improving temporal resolution for RNA conformational sensing. αHL_NetB also enhances the capture and threading of intrinsically disordered α-synuclein, demonstrating its versatility for protein sensing. Together, these results validate β-barrel domain engineering as a powerful strategy for tuning nanopore function and position the αHL_NetB as a versatile and promising platform for next-generation single-molecule biosensing and sequencing technologies.

## Methods & Materials

### Chimera protein expression and purification

The wild-type αHL and chimera proteins were expressed and purified following previously established protocols. In short, verified chimera proteins plasmids were transformed into competent E. coli BL21(DE3) pLysS cells (Sigma). A single colony from an overnight culture grown on ampicillin-containing agar plates (100 µg/mL) at 37 °C was inoculated into 100 mL of LB medium supplemented with ampicillin (50 µg/mL) and incubated at 37 °C with shaking at 160 rpm until an OD_600_ of 0.7 was reached. Protein expression was then induced overnight at 18 °C by adding 1 mM IPTG. Cells were harvested by centrifugation at 3800 rpm and incubated at 4 °C for 20 minutes before lysis in a buffer containing 50 mM Tris-HCl (pH 8.0), 0.5 M NaCl, 0.1% Triton X-100, and 10 mM imidazole. To facilitate lysis, 5 mM MgCl₂, 1 mg/mL lysozyme, and 100 units of Benzonase Nuclease (Cat. No.: E1014) were added, followed by sonication and centrifugation at 29,000 × g for 45 minutes to clear cellular debris. Purification of wild-type αHL and chimera proteins was carried out using immobilized metal-affinity chromatography (IMAC) with a TALON (Cobalt) resin. After binding, proteins were eluted with a buffer containing 50 mM Tris-HCl (pH 8.0), 0.5 M NaCl, 0.38 mM DDM, and 250 mM imidazole. Purity and oligomeric state (monomeric and heptameric forms) were assessed by SDS-PAGE (12%), and protein concentration was determined using Pierce^TM^ BCA Protein Assay Kit (23227, Thermo Scientific).

### α-synuclein expression and purification

Wild-type α-synuclein (1–140) was expressed in *E. coli* BL21(D3) and purified as previously described (42–44). In brief, protein expression was induced with IPTG at a final concentration of 1 mM and cultures were incubated overnight at 21°C in LB medium. Cells were harvested and the pellet was lysed by heat treatment followed by ultrasonication. The lysate was subjected to 1.75 M ammonium sulphate precipitation. The resulting pellet was resuspended in 25 mM Tris-HCl (pH 8.0, filtered) and loaded onto a HiTrap Q FF anion exchange chromatography column. The protein was eluted using a linear gradient of 25 mM Tris-HCl (pH 8.0), 800 mm NaCl. Fractions containing α-syn were pooled and precipitated again with ammonium sulphate (final concentration 1.75 mM). The pellet was resuspended in 6 M Guanidine Hydrochloride, 25 mM Tris-HCl (pH 7.4, filtered) and size exclusion chromatography was performed using a HiLoad 16/600 SuperDex 75 pg column. After dialysis against 800 mM ammonium bicarbonate, the protein concentration was determined by measuring UV absorbance at 280 nm using an extinction coefficient of 5960 M⁻¹cm⁻¹, followed by aliquoting and lyophilization.

### Hemolysis assay

MBSA buffer (150 mM NaCl, 10 mM MOPS, pH 7.4, supplemented with 0.1% BSA) was prepared for washing rabbit red blood cells (rRBCs). The raw rabbit blood cells in EDTA-K2 solution were purchased from Envigo. rRBCs were washed repeatedly on ice and then centrifuged at 1400 rpm for 4 min until the supernatant became clear. Then, the supernatant was removed, and MBSA buffer was added to prepare a 10% rRBCs solution. Afterwards, the 10% rRBCs were diluted to 1% rRBCs for the hemolysis assay.

Wild-type αHL and chimera proteins, dissolved in a solution containing 50 mM Tris-HCl (pH 8.0), 0.5 M NaCl, 250 mM imidazole, and 0.38 mM DDM, were utilized for the hemolysis assay. The initial concentrations of wild-type αHL and chimera proteins are as follows: αHL, 0.26 mg/mL; αHL_Aerolysin, 0.32 mg/mL; αHL_Anthrax, 0.35 mg/mL; αHL_CEL-III, 0.35 mg/mL; αHL_NetB, 0.33 mg/mL; αHL_Epsilon, 0.31 mg/mL; and αHL_VCC, 0.35 mg/mL. 2 µL of them were mixed with 98 µL MBSA buffer. Next, 50 μL was transferred to the second well and mixed with 50 μL of MBSA buffer for dilution. This process was repeated sequentially through the 12th well, resulting in each well containing 50 μL of solution with serial two-fold dilutions.

Subsequently, 50 μL of fresh 1% rRBC suspension was added to each well. Hemolytic activities of wild-type αHL and chimera proteins were assessed using a non-treated, unsealed, flat-bottom 96-well plate (Greiner Bio-One™, Catalog No.: 655101) made of polystyrene, on a Synergy™ 4 (BioTek) plate reader. Light scattering absorbance at 595 nm was monitored over 2 hours at room temperature to measure the decrease associated with hemolysis. Three separate experiments were performed independently.

### Single-channel current measurements and data analysis

A lipid bilayer was formed by introducing 10 mg/mL of 1,2-diphytanoyl-sn-glycero-3-phosphocholine (DPhPC) or 20 mg/ml 1,2-Dioleoyl-*sn*-glycerol-3-phosphocholine (DOPC): 1,2-dioleoyl-sn-glycero-3-phospho-(1’-rac-glycerol) (DOPG) = 4:1 into a solution containing 1 M KCl and 10 mM HEPES, adjusted to pH 7.4. The bilayer spanned an approximately 100 µm diameter aperture in a Teflon film, separating the setup into cis and trans chambers, each containing 600 µL of the solution. Wild-type αHL or chimera proteins were prepared by diluting a 0.1 mg/mL stock solution 1000-fold with Millipore water. To form a single nanopore within the lipid bilayer, 0.2–0.5 µL of the diluted protein solution was added to the cis chamber. After one pore formation, ssDNA was introduced to the *cis* side to a final concentration of 1 μM. PreQ1 riboswitch aptamers were introduced into the *cis* chamber at a final concentration of 200 nM. The aptamers, dissolved in sterile Millipore water, were refolded by heating them at 80°C for 5 minutes in a refolding buffer (50 mM Hepes-KOH, pH 7.5, 100 mM KCl, 4 mM MgCl₂), then slowly cooled down to room temperature. The final concentration of PreQ1 was 20 μM. α-syn was added into the *cis* chamber at a final concentration of 1 nM. Ionic currents across the nanopore were recorded using an Axopatch 200B amplifier (Molecular Devices), low-pass filtered at 1 kHz, and digitized with a DigiData 1440A converter (Molecular Devices) at a sampling rate of 250 kHz. Nanopore current traces of ssDNA and preQ1 riboswitch aptamers were analyzed with Clampfit 10.7 software (Molecular Devices). Residual currents (I_r_/I_0_ (%)) were obtained by fitting event current amplitude histograms using Gaussian fitting in Origin software, while the most probable event durations were determined by fitting event duration histograms similarly. Event frequency was calculated by dividing the number of ssDNA translocation events by the total recording time. The ssDNAs were synthesized from Microsynth. Standard deviations were determined from at least three independent experiments. Nanopore current traces of α-syn, recorded at an applied voltage of +100 mV, were analyzed using Python to extract residual current (I_r_/I_0_ (%)) and event duration values for single translocation events. Event frequency was calculated as the reciprocal of the interevent interval (1/τ_IE_), where τ_IE_ is the time between successive translocation events. All nanopore sensing experiments were performed at room temperature.

### Computational simulation

#### MD Simulation

Chimera pores were predicted by MODELLER All the MD runs were carried out using NAMD 2.14(45) with a time step Δt = 2.0 fs (unless otherwise stated) and particle mesh Ewald(46) method with a 1.0 Å spaced grid for long-range electrostatic interactions. A cutoff of 12 Å (switching distance of 10 Å) was used for the shortrange nonbonded interactions, while using 1-4 scaling. This scaling parameter was set to 1.0. Periodic boundary conditions with an orthogonal box are used. A Langevin thermostat was used for all the simulations while a Nosé-Hoover Langevin piston pressure control was employed for constant pressure simulations(47). CUFIX corrections(48) for ions were applied for CHARMM36(49) simulations.

#### Currents

A water box (150Å × 150Å × 190Å) used for conductivity measurements is generated by VMD solvate package(50) while K^+^ and Cl^−^ ions are added using the VMD autoionize package. The resulting pdb and psf are employed for the simulation using the CHARMM36 force field(51) with TIP3P(52) water model. After 5000 steps of energy minimization, the systems is equilibrated by a 0.1 ns NPT run (Δt = 1.0 fs). Non-equilibrium simulations were run with an external electrical field E = (0, 0, *E_z_*) with

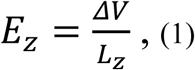

where *ΔV* is the applied voltage and *L_z_* is the size of the box along the z axis. A thermostat is applied to the oxygen atoms of the water. For each *ΔV*, we run 50 ns, the first 10 ns being discarded in the analysis. The electric current *I* is measured as it follows(53,54). During production run, trajectories were printed every τ = 20 ps. In post-processing, we calculated(53,54).

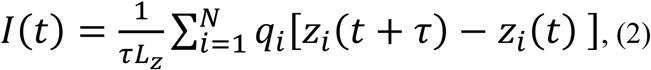

where *I(t)* is the average current in the interval t ∈ [t, t+τ], q_i_ is the charge of the *i−th*, *z_i_* its z-coordinate, and *N* is the number of ions in the system. *I* is estimated by averaging the current time trace from Eq. (2), while errors are calculated using a block average protocol, with a block size 10 ns. Cations and anions’ contributions to *I* are calculated, restricting the summation in Eq. (2) only to K^+^ and Cl^−^.

#### Nanopore and membrane set-up and equilibration

The protocol used to build and to equilibrate the system is similar to the one reported in Di Muccio et al.(54) that is also similar to the one used in previous works(33,55). The pore axis is aligned with the z-axis of the reference system. The pore is inserted in a POPC bilayer membrane generated using VMD.6 Lipid molecules overlapping with the nanopore or located in the pore lumen are removed. Subsequently, water and ions are added using the solvate and ionize VMD plugins. (6) After 5000 steps of energy minimization, a first 494 ps NPT equilibration step (P = 1 atm, flexible cell, constant ratio, T = 300 K, time step Δt = 0.5 fs) is performed. In this first step, external forces are applied to the water molecules to avoid their penetration into the membrane. The *C_α_* atoms of the nanopore are kept fixed and the phosphorus atoms of the lipid heads are harmonically constrained to their z-positions (spring constant k = 1 kcal/mol·Å^2^). An initial temperature ramp is imposed (every 5 ps the velocities are rescaled to a temperature of 0, 25, 50, 75, 100, 125, 150, 175, 200, 225, 250, 275, 300 K) for a smooth relaxation of the system. In essence, in this step, the lipid tails and the electrolyte relax while the nanopore backbone is fixed. In a second, 1 ns long (Δt = 1 fs) NPT equilibration step, all the constraints on the membrane and the protein are removed. All the other control settings are as in the previous step. The last equilibration step is an NPT run of 1 ns (time step 1 fs) with no constraints on the lipids and no external forces to keep the water molecules out of the membrane. Subsequently, non-equilibrium simulations were run with an external electrical field E = (0, 0, *E_z_*), Eq. (1). Nanopore *C_α_* are constrained while the phosphorus atoms of the lipids are fixed. A thermostat is applied to protein and lipids with the exception of the constrained atoms. For each non-equilibrium simulation we run 40 ns, the first 10 ns being discarded from the analysis. Trajectories are printed every τ = 20 ps and we calculated currents using the same protocol employed for the triperiodic water box, see Eq. (2). The EOF is measured similarly, computing the summation over the oxygen atoms of the water molecules and just counting the molecules. For each *ΔV*, we run different replicas of the above simulation protocol (including the equilibration). For each replica, average currents and EOF are calculated. The final estimation of the current is obtained by averaging the results for different replicas, while its error is estimated dividing the standard deviation of the averages by the square root of number of replicas. Charge density maps were computed as in Di Muccio et al (54).

## Author Contributions

J.L. conceived the experiments, C.L. and E.K. conducted the experiments, M.R. ran the molecular simulations, C.L., E.K. and M.R. analyzed the results, C.L. wrote the manuscript, and all authors revised the manuscript.

## Acknowledgements

The authors acknowledge funding from the Swiss National Science Foundation (310030-197626 & 10002967 to J.L.), the BrightFocus Foundation (A20201759S to J.L.), the China Scholarship Council (202106260035 to C.L.). Molecular Dynamics Simulations were run using HPC computational resources provided by Swiss National Supercomputing Centre (CSCS), project ID lp33.

