## Supplemental information for "β-Barrel domain swapping in α-hemolysin enables enhanced single-molecule biomolecule sensing"

**Supporting information**

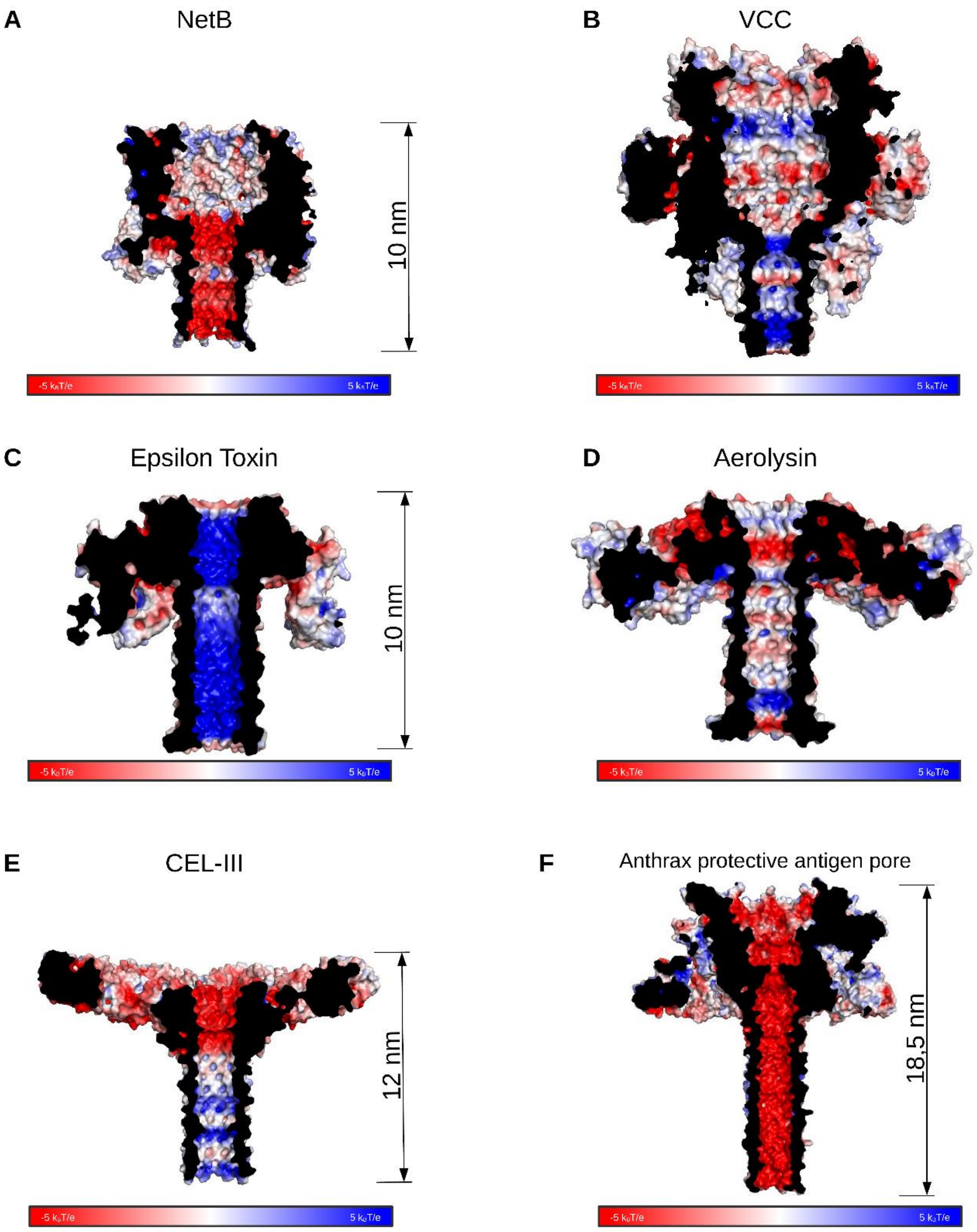


**Figure S1. Electrostatic surface potential maps of β-barrel pore-forming toxins.** Electrostatic surface representations of six representative β-pore-forming proteins were generated to visualize charge distribution within the barrel lumen and surrounding regions. Each panel shows a cross-sectional view of the electrostatic potential, colored from red (negative potential) to blue (positive potential), generated using APBS (1) plugin for PyMOL software (Schrodinger, LLC (2015) The PyMOL Molecular Graphics System, Version 2.3.0.). (A) NetB (PDB ID: 4H56 (2)), (B) VCC (PDB ID: 3O44 (3)), (C) Epsilon toxin (PDB ID: 6RB9 (4)), (D) Aerolysin (PDB ID: 5JZT (5)), (E) CEL-III PDB ID: 3W9T (6)), (F) Anthrax protective antigen pore (PDB ID: 6UZB (7)). The electrostatic potential scales (kbT/ec) are shown below each structure. These visualizations highlight distinct charge distributions within the luminal surfaces of the pores, which may influence ion selectivity and molecular translocation properties.

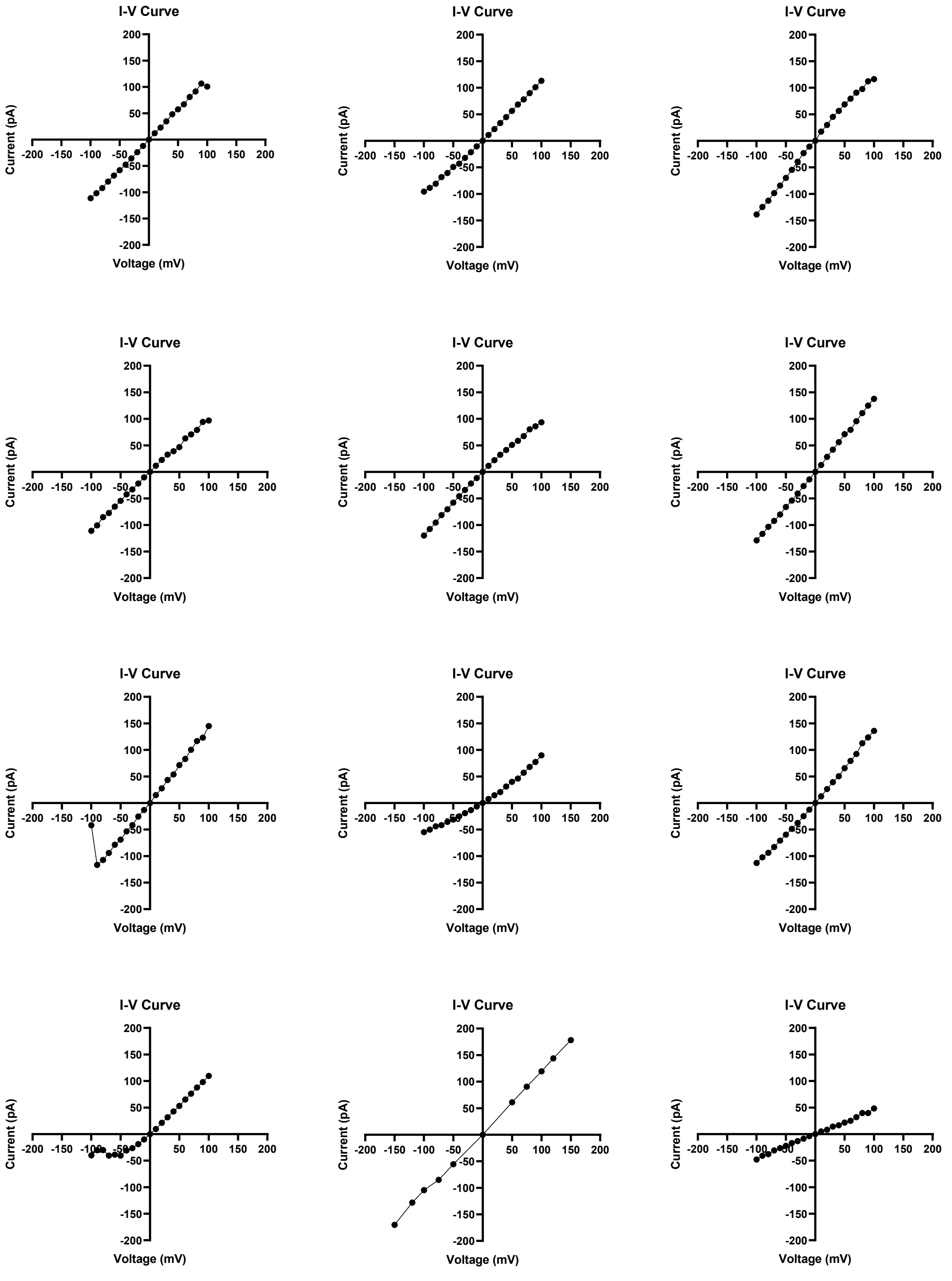


**Figure S2. Current–voltage (I–V) characteristics of independently reconstituted αHL_NetB chimera nanopores.** I–V curves recorded from individual αHL_NetB chimera nanopores reconstituted into planar lipid bilayers under symmetrical buffer conditions (10 mM HEPES, 1 M KCl, pH 7.4). Each trace corresponds to a different pore obtained from independent bilayer experiments. The pores exhibit measurable ionic conductance across the tested voltage range; however, noticeable gating behavior and current instability are observed, particularly at higher applied potentials. These features are reflected by current fluctuations and deviations from ideal linear ohmic behavior, indicating reduced stability of the chimera compared to more stable pore constructs. The variability between traces further highlights pore-to-pore differences in conductance and stability.

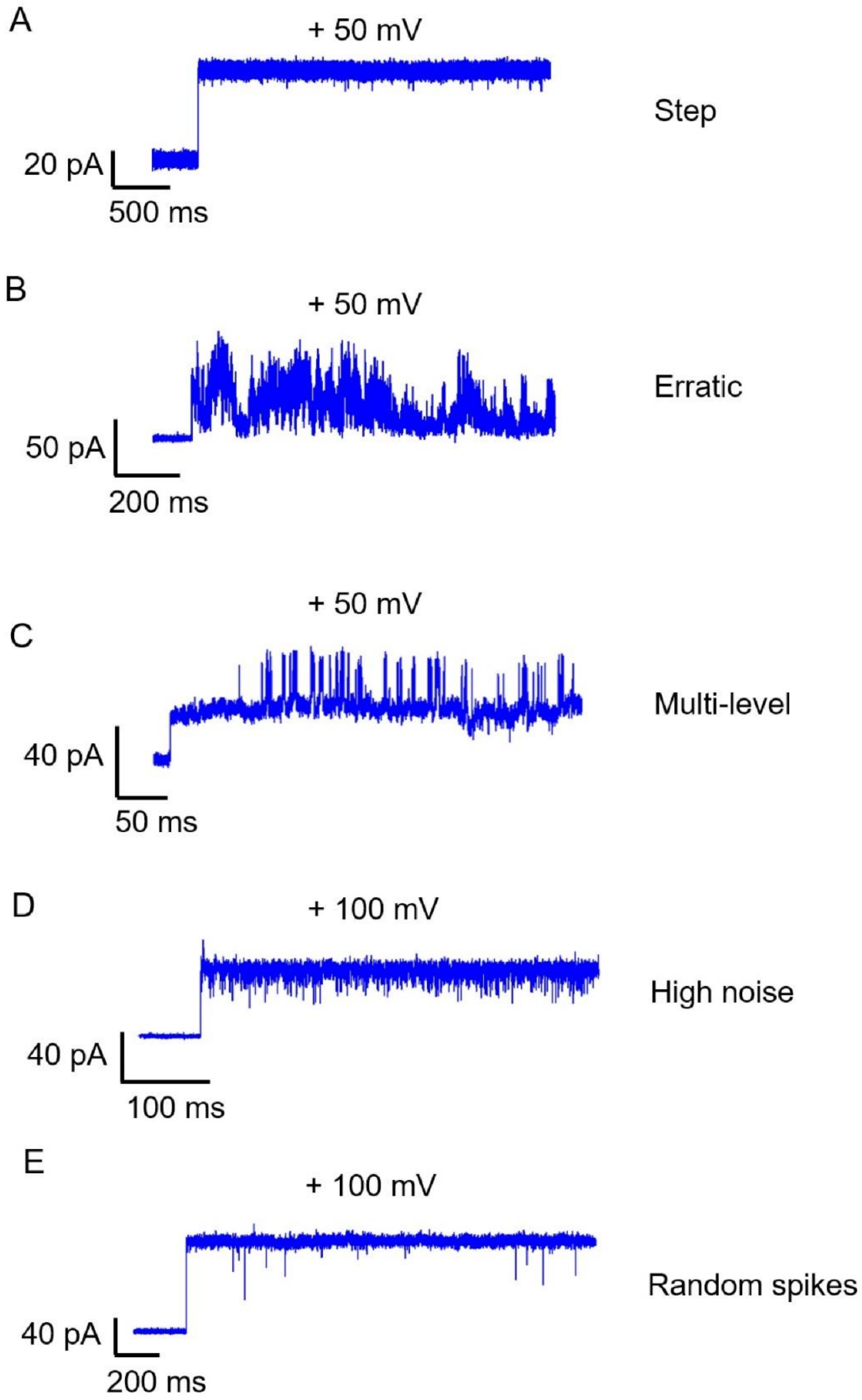


**Figure S3. Representative electrical recordings illustrating the operational definitions of stable and unstable single-pore formations.** Representative current traces showing examples of different levels of pore stability. (A) A stepwise current transition followed by a steady baseline with minimal fluctuation is defined as a stable single-pore formation and is suitable for nanopore sensing measurements. In contrast, traces classified as unstable include (B) erratic current behavior without a well-defined open state, (C) multi-level conductance states indicating frequent transitions between substates, (D) excessive baseline noise, and (E) random current spikes. These representative examples establish the criteria used in this study to distinguish stable pores from unstable pore insertions that are not appropriate for reliable single-nanopore sensing experiments. The pore stability analysis shown in Figures 3 and 4 was based on calculating the percentage of stable pore formation (A) and unstable pore formation (B–E) across independent pore formation experiments.

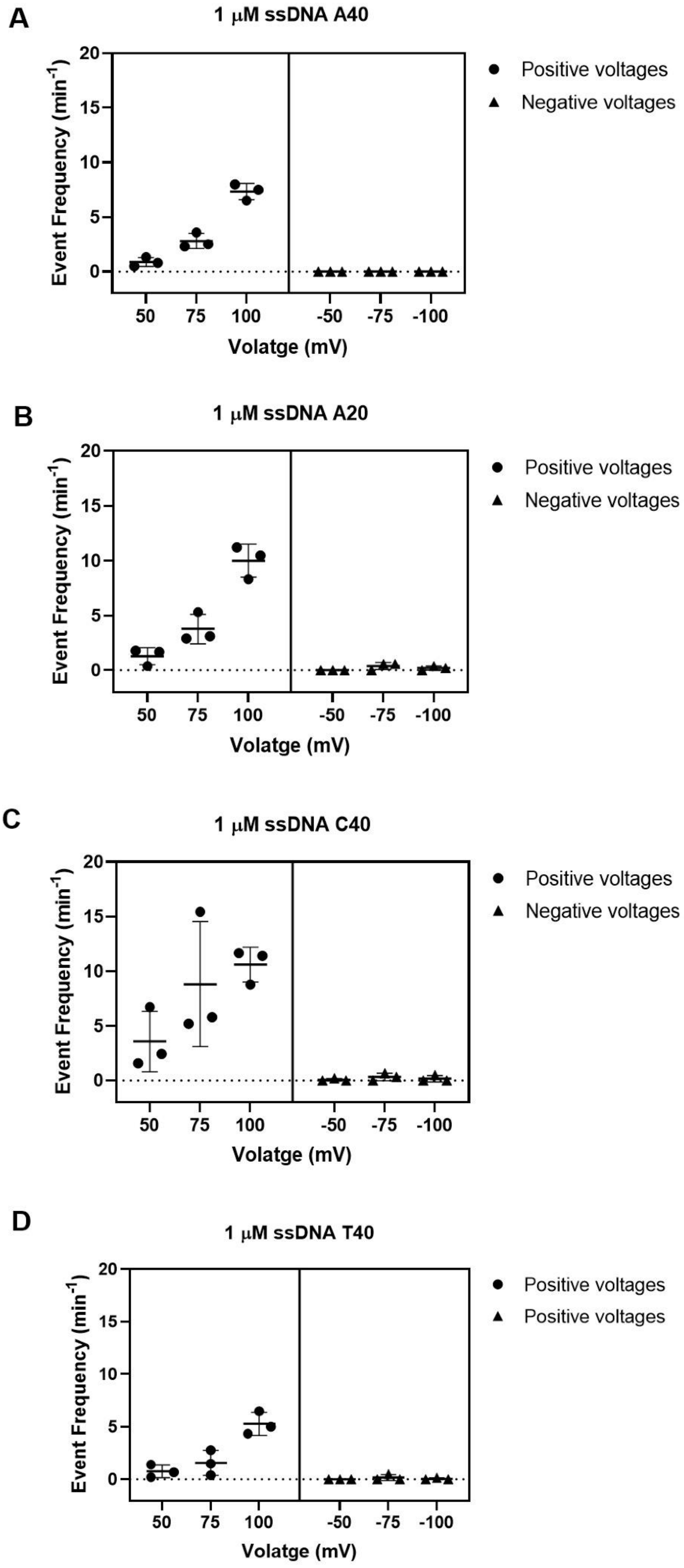
A
1 μM ssDNA A40
Event Frequency (min-1)
20
15
10
5
0
50
75
100
-50
-75
-100
Volatge (mV)
Positive voltages
Negative voltages
B
1 μM ssDNA A20
Event Frequency (min-1)
20
15
10
5
0
50
75
100
-50
-75
-100
Volatge (mV)
Positive voltages
Negative voltages
C
1 μM ssDNA C40
Event Frequency (min-1)
20
15
10
5
0
50
75
100
-50
-75
-100
Volatge (mV)
Positive voltages
Negative voltages
D
1 μM ssDNA T40
Event Frequency (min-1)
20
15
10
5
0
50
75
100
-50
-75
-100
Volatge (mV)
Positive voltages
Positive voltages

**Figure S4. Voltage-dependent event frequency of different homopolymeric ssDNA oligonucleotides recorded using the αHL_NetB nanopore.** Event frequencies were measured for 1 μM concentrations of four different ssDNA homopolymers: (A) A40, (B) A20, (C) C40, and (D) T40. Frequencies were recorded at various applied voltages ranging from +50 mV to +100 mV and −50 mV to −100 mV. Each data point represents the mean event frequency (events per minute) from independent measurements, with error bars indicating standard deviation. Circles (●) represent measurements at positive voltages, while triangles (▲) correspond to negative voltages. All homopolymers showed a strong bias for event detection at positive voltages, with negligible activity observed at negative voltages. Buffer conditions: 10 mM Hepes, 1 M KCl, pH 7.4. Error bars represent standard deviations from independent replicates.
